# Parallels between experimental and natural evolution of legume symbionts

**DOI:** 10.1101/250159

**Authors:** Camille Clerissi, Marie Touchon, Delphine Capela, Mingxing Tang, Stephane Cruveiller, Matthew A. Parker, Lionel Moulin, Catherine Masson-Boivin, Eduardo P.C. Rocha

## Abstract

The emergence of symbiotic interactions has been studied using population genomics in nature and experimental evolution in the laboratory, but the parallels between these processes remain unknown. We compared the emergence of rhizobia after the horizontal transfer of a symbiotic plasmid in natural populations of *Cupriavidus taiwanensis*, over 10 MY ago, with the experimental evolution of symbiotic *Ralstonia solanacearum* for a few hundred generations. In spite of major differences in terms of time-span, environment, genetic background and phenotypic achievement, both processes resulted in rapid diversification dominated by purifying selection concomitant with acquisition of positively selected mutations. The latter were lacking in the plasmid carrying the genes responsible for the ecological transition. Instead, adaptation targeted the same set of genes leading to the cooption of the same quorum-sensing system. Our results provide evidence for similarities in experimental and natural evolutionary transitions and highlight the potential of comparisons between both processes to understand symbiogenesis.

---

Biological adaptations have traditionally been evaluated by inferring the evolutionary history of organisms from the genomic, morphological, and phenotypic comparison of natural isolates, including fossil records when they were available. Recently, these approaches have been increasingly complemented by experimental evolution studies. The latter can be done on controlled environments and provide nearly complete “fossil” records of past events because individuals from intermediate points in the experiment can be kept for later analysis ^1,2^. Sequencing and phenotyping of evolved clones provides crucial information on the mechanisms driving adaptation in simplified environments. Yet, there is little data on the adaptation of lineages in the case of complex adaptations requiring numerous steps and even less on how they recapitulate natural processes (but see ^3,4^), raising doubts on the applicability and relevance of experimental evolution studies to understand natural history ^5^.

Adaptations to new and complex environments, such as ecological transitions towards pathogenic or mutualistic symbiosis, are often initiated by the acquisition via horizontal transfer of genes that provide novel functionalities ^6^. For example, the extreme virulence of *Shigella* spp., *Yersinia pestis*, or *Bacillus anthracis* results from the acquisition of plasmid-encoded virulence factors by otherwise poorly virulent clones. These novel genetic systems often require subsequent regulatory rewiring, a process that may take hundreds to millions of years *in natura* ^7^. A striking case of horizontal gene transfer (HGT)-mediated transition towards mutualism concerns the rhizobium-legume symbiosis, a symbiosis of major ecological importance that contributes to ca. 25% of the global nitrogen cycling. Rhizobia induce the formation of new organs, the nodules, on the root of legumes, which they colonize intracellularly and in which they fix nitrogen to the benefit of the plant ^8^. These symbiotic capacities emerged several times in the natural history of α- and β-Proteobacteria, from the horizontal transfer of the key symbiotic genes into soil free-living bacteria (*i.e.*, the *nod* genes for organ formation and the *nif/fix* genes for nitrogen fixation) ^9–11^. This process resulted in hundreds of rhizobial species scattered in 14 known genera, including the genus *Cupriavidus* in β-proteobacteria ^12^.

Transition towards legume symbiosis has recently been tested at the laboratory time-scale using an experimental system ^13^. A plant pathogen was evolved to become a legume symbiont by mimicking the natural evolution of rhizobia at an accelerated pace. First, the plasmid pRalta^LMG19424^ - encoding the key genes allowing the symbiosis between *C. taiwanensis* LMG19424 ^14^ and *Mimosa* – was introduced into *Ralstonia solanacearum* GMI1000. The resulting chimera was further evolved under *Mimosa pudica* selective pressure. The chimeric ancestor, which was strictly extracellular and pathogenic on *Arabidopsis thaliana* - but not on *M. pudica* and unable to nodulate it - progressively adapted to become a legume symbiont during serial cycles of inoculation to the plant and subsequent re-isolation from nodules ^13,15,16^. Several adaptive mutations driving acquisition and/or drastic improvement of nodulation and infection were previously identified ^13,17,18^. Lab-evolution was accelerated by stress-responsive error-prone DNA polymerases encoded in the plasmid that increased the mutation load *ex planta* ^19^.

Here we compare the natural and experimental evolutions of *Mimosa* symbionts in the *Cupriavidus/Ralstonia* branch using population genomics and functional enrichment analyses. We traced the natural evolutionary history of *Cupriavidus taiwanensis* and provide evidence that, despite significant differences in terms of time frame, protagonists, and environmental context, there were very significant parallels in the two processes.

## Results

### Diversification of naturally and experimentally evolved *Mimosa* symbionts

We previously generated 18 independent symbiotic lineages of the *R. solanacearum* GMI1000-pRalta^LMG19424^ chimeras that nodulate *M. pudica* ^15^. Each lineage was subject to 16 successive cycles of evolution in presence of the plant. We isolated one clone in each of the lineages after the final cycle to identify its genetic and phenotypic differences relative to the ancestor. The symbiotic performances of the evolved clones improved in the experiment with wide variations between lineages. Some clones were able to produce nodules massively and intracellularly infected (Fig. 1A). Yet none of them fixed nitrogen to the benefit of the plant at this stage. In addition to a total of ca. 1200 point mutations relative to the ancestral clones ^15^, we detected several large deletions in all clones (Fig. 1A). The positions of point mutations were different between lineages, but some genes and many functional categories were affected in parallel (Fig. 1B). In contrast, the deletions showed frequent parallelisms at the nucleotide level. They occurred in homologous regions of the symbiotic plasmid and were systematically flanked by transposable elements that probably mediated their loss by recombination (Table S1).

**Figure 1.**
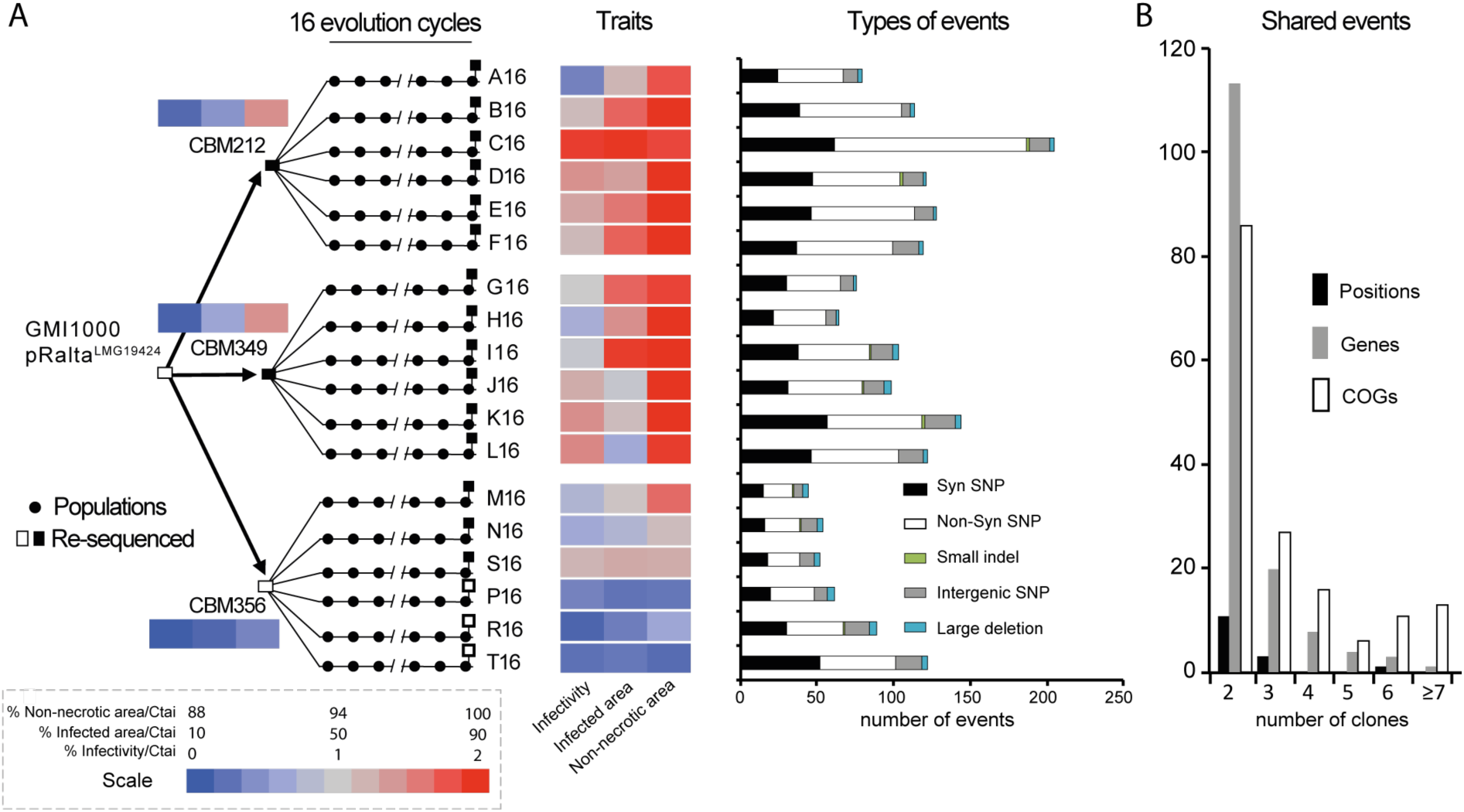
Experimental evolution of *Ralstonia* and associated symbiotic and genomic changes. **A**.An ancestor chimeric clone evolved to give origin to three clones able to nodulate *M. pudica*. Each of these clones was then evolved in 18 independent lineages using 16 serial nodulation cycles. This process led to improved infectivity (number of viable bacteria recovered per nodule) and intracellularly-infected area per nodule section and a decrease of necrotic area per nodule section (heatmap on traits). Except clones CBM356, P16, R16 and T16 (white squares), all acquired the ability of intracellular infection (black squares.) The events identified at the end of the 16 evolution cycles for each lineage are indicated on the right (see list of deletions in Table S1 and other mutations in Table S12). **B**. Number of shared events between lineages, *i.e.* the number of positions, genes, and COG categories of genes that were mutated in two or more lineages.

We sequenced, or collected from public databanks, the genomes of 58 *Cupriavidus* strains to study the genetic changes associated with the natural emergence of *Mimosa* symbionts in the genus and to compare them with those observed in the experiment (see supplementary Text S1 and associated tables for data sources, coverage, and details of the results). The phylogeny of the genus core genome was well resolved, showing that 44 out of the 46 genomes with the *nod* and *nif* genes were in the monophyletic *C. taiwanensis* clade (Fig. 2). The two exceptions, strains UYPR2.512 and amp6, were placed afar from this clade in the phylogenetic tree and are clearly distinct species. *C. taiwanensis* strains are *bona fide* symbionts since they fixed nitrogen in symbiosis with *M. pudica* ^20,21^. Unexpectedly, the average nucleotide identity (ANIb) values between *C. taiwanensis* strains were often lower than 94%, showing the existence of abundant polymorphism and suggesting that *C. taiwanensis* is not a single species, but a complex of several closely related ones (Fig. 2 and S1, Text S1, Table S2). Together, *C. taiwanensis* strains had a core genome of 3568 protein families and an open pan genome, 3.4 times larger than the average genome. Hence, this complex of species has very diverse gene repertoires.

**Figure 2.**
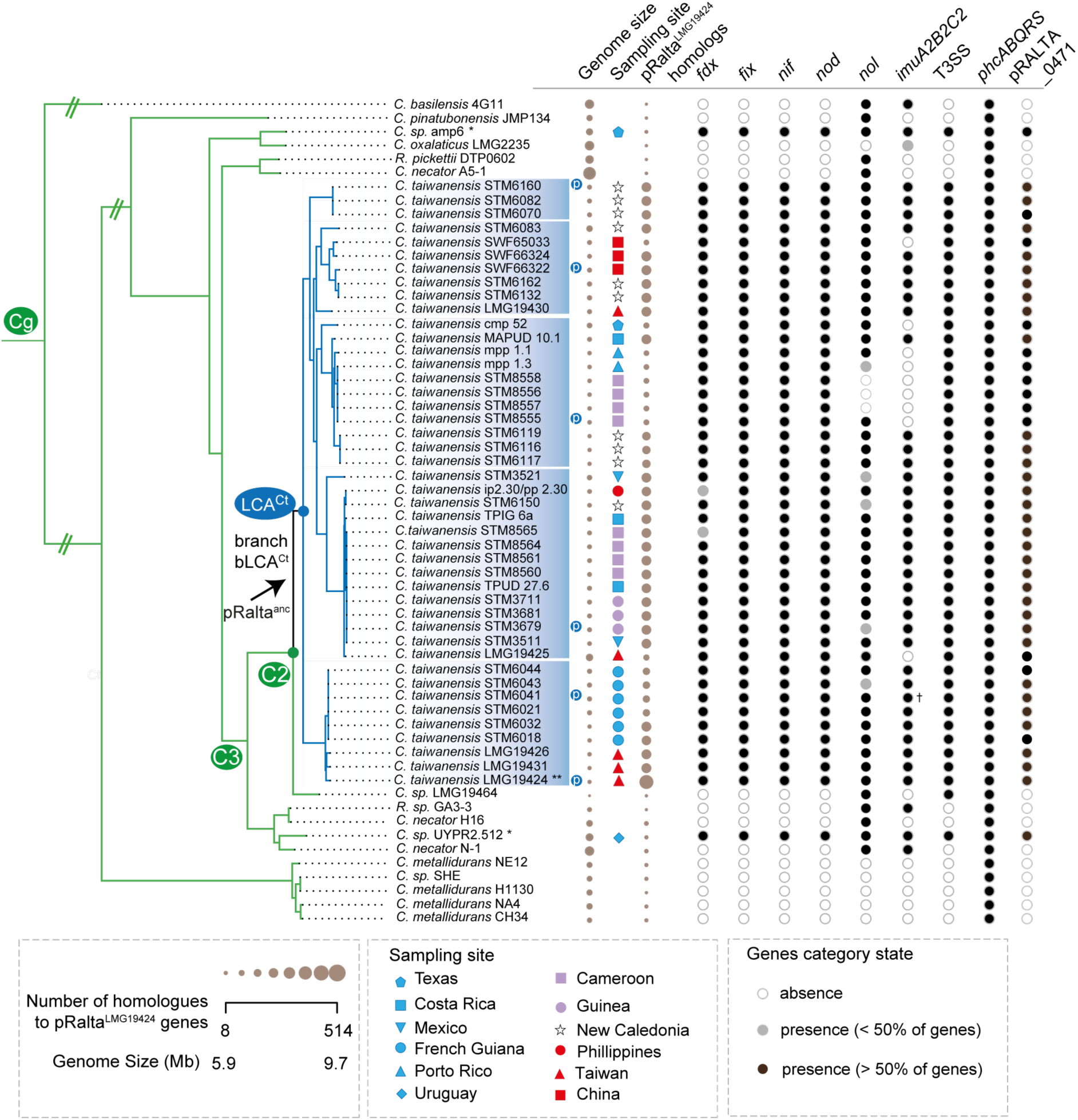
Distribution of symbiotic genes, the mutagenic cassette, T3SS, *imuA2B2C2* and *phcABQRS* within the 60 strains of *Cupriavidus*. See Fig. S1 for the complete tree of the genus *Cupriavidus* and *Ralstonia* without simplifications in branch length. The arrow indicates the most parsimonious scenarios for the acquisition of pRalta (inferred using the MPR function of the ape package in R). This is the branch before the LCA^Ct^. The node LCA^Ct^ indicates the last common ancestor of *C. taiwanensis*. Circles indicate absence (white), presence of less than 50% of the genes (light grey) and presence of more than 50% of the genes (black). Note that most rhizobia possess the pRalta_0471 gene which is located downstream a *nod* box in LMG19424. The size of the circles for *Genome size* and *pRalta homologs* is proportional to the value of the variable. Sampling sites are coded according to geographic origins. Clusters were computed according to different thresholds of ANIb (as indicated in the text and in Figs. S1 and S8). Symbols: Ct, C2, C3 and Cg: LCA of clades analyzed in this study. p (in a blue circle): plasmid re-sequenced by PacBio. *: two rhizobia are not part of *C. taiwanensis*. **: *C. taiwanensis* reference strain used as pivot to compute searches of orthologs. † In the PacBio version of this genome *imuABC* is very similar to that of the reference strain, but is encoded in another plasmid.

### Parallel patterns of evolution upon the acquisition of the symbiotic plasmid

To compare the initial stages of adaptation in natural populations with those in experimental populations, we searched to identify when the rhizobial character (defined by the presence of the key symbiotic genes *nod* and *nif*/*fix*), was acquired in the genus *Cupriavidus* (Figs. 2 and S2). The most parsimonious reconstruction of the character in the phylogenetic tree revealed three independent transitions towards symbiosis: in the branch connecting the last common ancestor of *C. taiwanensis* and its immediate ancestor (branch before LCA^Ct^, hereafter named bLCA^Ct^), and in the terminal branches leading to strains UYPR2.512 and amp6. In agreement with these conclusions, we found very few homologs of the 514 pRalta^LMG19424^ genes in the genomes of UYPR2.512 (8.3 %) or amp6 (6.4%) once the 32 symbiotic genes were excluded from the analysis. These few homologs in the plasmid also showed significantly lower values of sequence similarity than the core genes of the genus (p <0.01, Wilcoxon test). We then used birth-death models to identify the acquisitions of genes in the branch bLCA^Ct^ (Fig. 2, Table S3). This analysis highlighted a set of 435 gene acquisitions that were present in pRalta^LMG19424^, over-representing functions such as symbiosis, plasmid biology, and type 4 secretion system (Table S4). These results are consistent with a single initial acquisition of the plasmid in this clade. PacBio resequencing of five strains representative of the main lineages, putative novel species, of *C. taiwanensis* confirmed the ubiquitous presence of a variant of pRalta encoding the symbiotic genes (Table S5). Finally, while most individual *C. taiwanensis* core gene trees showed some level of incongruence with the concatenate core genome tree, an indication of recombination, this frequency was actually lower in the core genes of the plasmid (p <0.04, Fisher’s exact test). Similarly, there were fewer signals of intragenic recombination in plasmid core genes (PHI, p <0.001, same test). This suggests that the plasmid inheritance was mostly vertical within *C. taiwanensis*. We thus concluded that the three rhizobial clades evolved independently and that the acquisition of the ancestral symbiotic plasmid of *C. taiwanensis* should be placed at the branch bLCA^Ct^. The date of plasmid acquisition was estimated using a 16S rRNA clock in the range 12-16 MY ago. Although these dating procedures are only approximate, the values are consistent with the low ANIb values within *C. taiwanensis* and are posterior to the radiation of its most typical host (*Mimosa* ^22^).

Since the experiment only reproduced the initial stages of symbiogenesis, parallels between experimental and natural adaptation should be most striking at the branch bLCA^Ct^, *i.e.*, during the onset of natural evolution towards symbiosis. The evolution experiment showed transient hypermutagenesis caused by the expression of the *imuA2B2C2* plasmid cassette *ex planta* ^19^. The long timespan since the acquisition of the plasmid precluded the analysis of accelerated evolution in the branch bLCA^Ct^ (relative to others). Yet, we were able to identify the *imuA2B2C2* cassette in most extant strains, suggesting that they could have played a role in the symbiotic evolution of *Cupriavidus*. We then searched for genes with an excess of recombination or nucleotide diversity in the branch bLCA^Ct^, which revealed 90 recombining genes and 67 genes with an excess of genetic diversity in this branch relative to the *C. taiwanensis* sub-tree (Fig. S3 and Table S3). To identify the parallels between the experimental and natural processes, we identified the 2372 orthologs between the *R. solanacearum* and *C. taiwanensis* (Table S6), and added the 514 pRALTA genes in the chimera as orthologs. Clones of the evolution experiment accumulated significantly more mutations in genes whose orthologs had an excess of polymorphism at the onset of symbiosis in natural populations (P <0.001, Fisher’s test; Tables S7 and S8), revealing a first parallel between the natural and experimental processes. A second parallel was identified in the overall regimes of natural selection. Both the substitutions in the core genes of *C. taiwanensis* (Fig. S4), and the mutations observed in the experiment ^15^ showed an excess of synonymous changes relative to the expected ones given the number of non-synonymous mutations. This shows a predominance of purifying selection in both processes, in spite of the observed adaptation towards symbiosis.

### Adaptation in the genetic background, not in the symbiotic plasmid

The symbiotic plasmids carry many genes and induce a profound change in the lifestyle of the bacteria. We thus expected to identify changes in the plasmid reflecting its accommodation to the novel genetic background. The plasmid pRalta^LMG19424^ accumulated an excess of synonymous substitutions and a vast majority of the genetic deletions observed in the experiment (Fig. 3 and Table S9). Natural populations also showed more deletions in the plasmid, since from the 413 genes present in pRalta ^LMG19424^ and inferred to be present in LCA^Ct^ only 12% were in the core genome, which is 6 times less than found among the chromosomal genes present in *C. taiwanensis* LMG19424 and inferred to be present in LCA^Ct^ (p <0.001, Fisher’s exact test, Fig. 3B). The few pRalta ^LMG19424^ core genes are related to the symbiosis or to typical plasmid functions (conjugation) (Fig. 3). The rate of recombination could not be measured on the genomes from the experiments because it is undetectable at this level of sequence similarity between clones (which presumably makes it less important as driver of diversification). The few plasmid core genes show lower recombination rates (PHI and SH analyses,both p <0.01) than the chromosomal ones.

**Figure 3.**
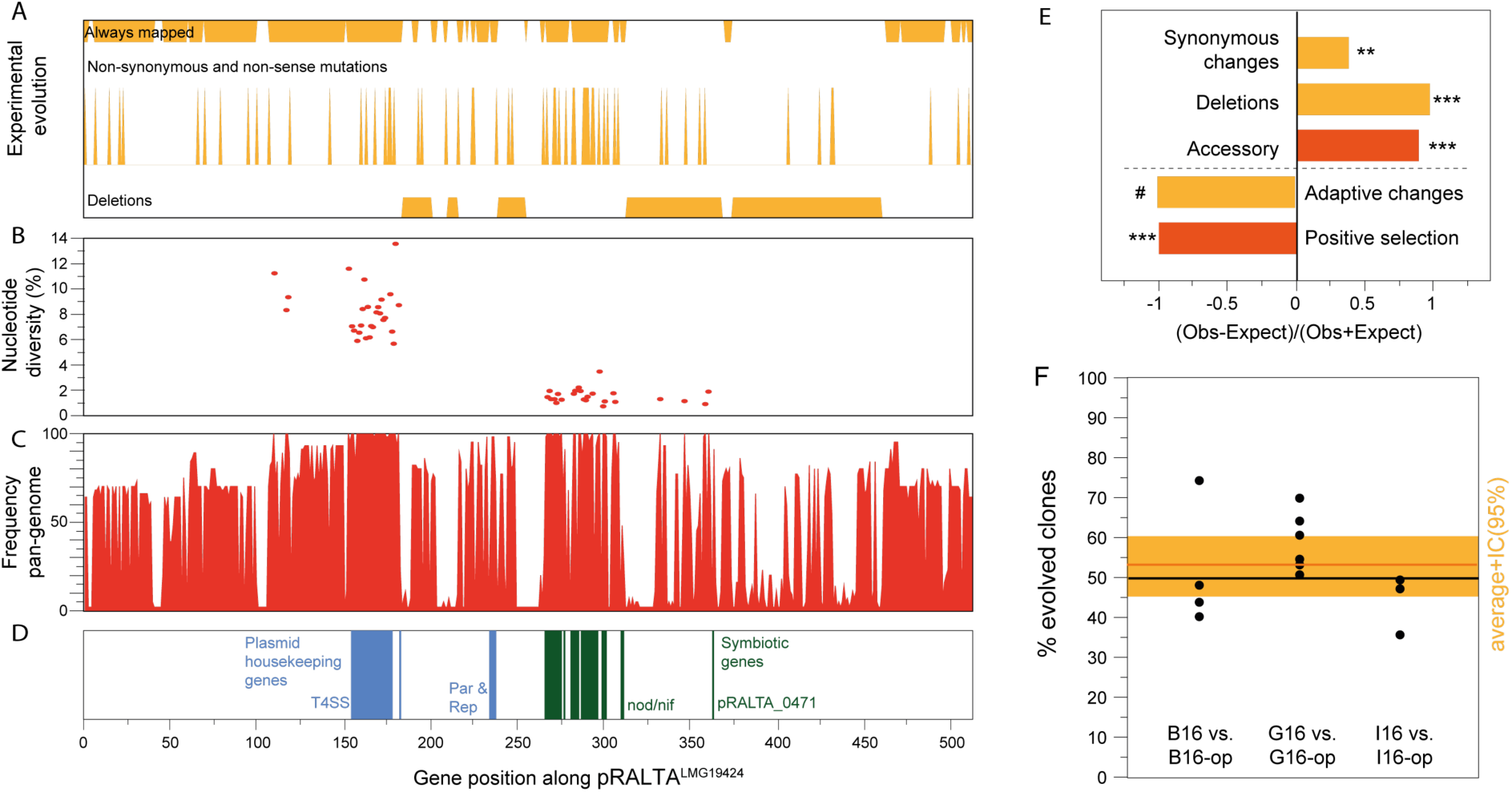
Analysis of the symbiotic plasmid of *Cupriavidus taiwanensis* LMG19424. **A**.Deletions, non-synonymous and non-sense mutations, and regions of the plasmid that could always be mapped to identify mutations in the experiment **B**. Nucleotide diversity of natural *C. taiwanensis* core genes: symbiotic genes accumulated much less diversity than the other genes. **C**. Frequency of each gene in the 44 *C. taiwanensis* (positional orthologs). **D**. Symbiotic and plasmid housekeeping genes. **E**. Observed over expected values for a number of traits in the plasmid natural (red) or experimental (orange) evolution (Tables S1, S3, and S8). **/*** significantly different from 1 (P < 0.01/0.001, Fisher’s exact tests for all but the test for “Synonymous changes” which was made by permutations, see Methods and Table S9). æ We could not find a single adaptive mutation in the plasmid in our previous works neither in the experiments in panel F. **F**. Impact of pRalta mutations on the *in planta* fitness of evolved clones. *M. pudica* plantlets were co-inoculated with pairs of strains at a 1:1 ratio and nodules were harvested at 21 dpi for bacteria counting. Each pair consisted of an evolved clone (B16, G16 or I16) and the same clone with the evolved pRalta replaced by the original one (B16-op, G16-op or I16-op). The orange horizontal bar represents the average and the large orange rectangle the 95% interval of confidence of the average (that includes the value 50% indicating that the two types of clones are not significantly different in terms of fitness).

To evaluate whether the observed rapid plasmid diversification was driving the adaptation to symbiosis *in natura*, we compared the rates of positive selection on plasmid and chromosomal genes in *C. taiwanensis*. We identified 325 genes under positive selection in the clade, and 46 specifically in the branch leading to LCA^Ct^ (analysis of 1869 and 1676 core genes lacking evidence of recombination using PHI, respectively, Table S3). Surprisingly, all 325 genes under positive selection were chromosomal (none was found among the core genes of the plasmid, Fig. 3E). In parallel, all mutations previously identified as adaptive in the evolution experiment were chromosomal ^13,17,18^. Since our previous analyses of mutations identified in the evolution experiment only focused on strongly adaptive genes, we evaluated the impact of pRalta^LMG19424^ mutations on the symbiotic evolution of *R. solanacearum* by replacing the evolved plasmid with the original pRalta^LMG19424^ in three evolved clones (B16, G16 and I16, thus generating strains B16-op, G16-op and I16-op, respectively). The relative *in planta* fitness of the new chimeras harboring the original plasmid were not significantly different from that of the experimentally evolved clones (Fig. 3F), showing that the adaptation of these strains did not involve mutations in the plasmid. Importantly, the original chimera had similar survival rates with and without the plasmid, suggesting that presence of the plasmid does not impact bacterial fitness in this respect (Tables S10 and S11). Although we cannot exclude that some events of positive selection in the plasmid may have passed undetected, nor that further symbiotic evolution of *R. solanacearum* will involve plasmid mutations, it appears that the genetic changes leading to improvement of the symbiotic traits mainly occurred in the chromosomes of *R. solanacearum* in the experiment, and of *C. taiwanensis* in nature, not on the plasmid carrying the symbiotic traits.

### Parallel co-option of regulatory circuits

We identified 436 genes with non-synonymous or non-sense mutations in the experiment (Table S12). This set of genes over-represented virulence factors of *R. solanacearum*, including the T3SS effectors, EPS production, and a set of genes regulating (*phcBQS*) or directly regulated (*prhI*, *hrpG,* and *xpsR*) by the central regulator PhcA of the cell density system that controls virulence and pathogenicity in *R. solanacearum* ^23^ (Fig. 4A and Table S13). Among them, mutations in the structural T3SS component *hrcV*, or in the virulence regulators *hrpG, prhI, vsrA*, and *efpR*, were demonstrated to be responsible for the acquisition or the drastic improvement of nodulation and/or infection ^13,17,18^.

**Figure 4.**
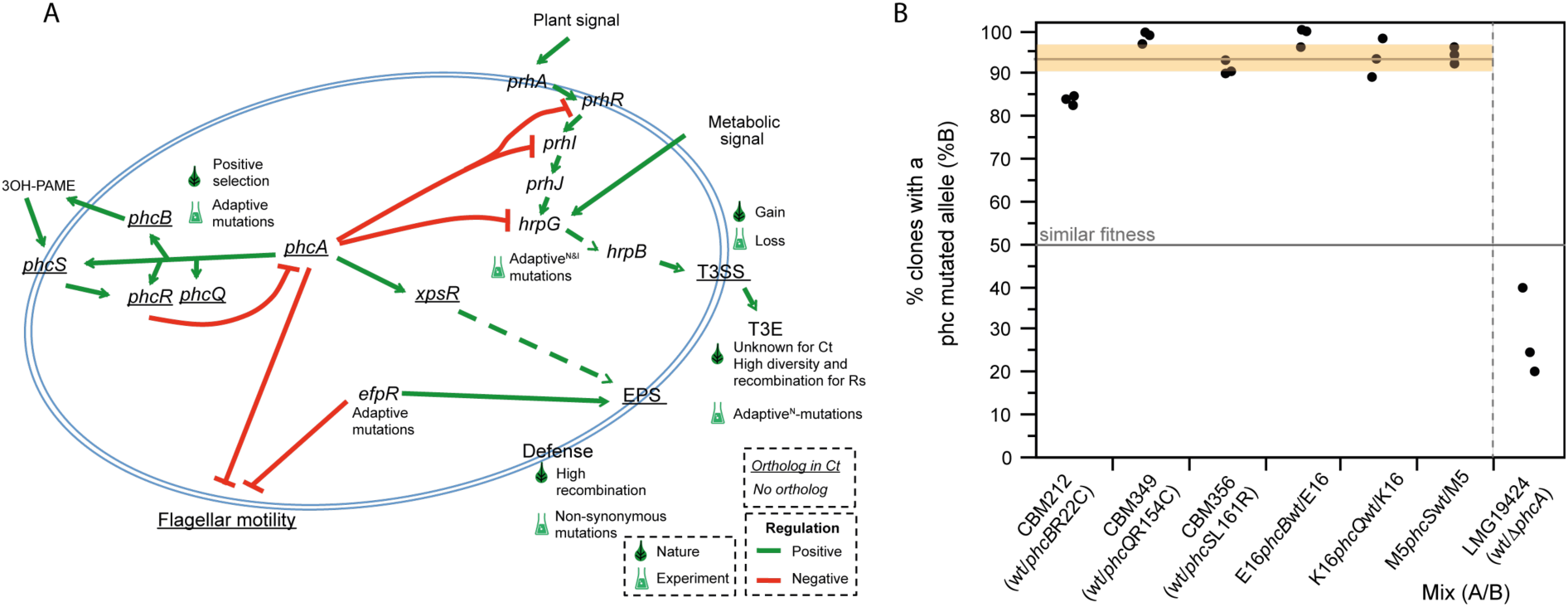
Virulence factors and regulatory pathways of *R. solanacearum* and their evolution in the evolution experiment. **A**. Schema of the major virulence factors and regulatory pathways mentioned in this study and their role in *R. solanacearum* (adapted from ^23^). Adaptive^N^ and adaptive^I^, represent the presence of adaptive mutations for nodulation and infection, respectively. Underline, genes or factors present in *C. taiwanensis*. The results of the enrichment analyses are in Tables S4, S13 and S21. **B**. Adaptive nature of the *phc* alleles evolved in the experiments and the recruitment of PhcA for symbiosis in the natural symbiont *C. taiwanensis* LMG19424. The horizontal grey line represents the average fitness of the evolved *phc* genes relative to the wild-type. The horizontal orange rectangle indicates the 95% interval of confidence for the mean. The results for *phc* are significantly different from the expected under the hypothesis that both variants are equally fit (horizontal line at 50%, p < 0.005, Wilcoxon test). The mean for the analysis of the mutant of PhcA (25%) is smaller than 50%, although the difference is at the edge of statistical significance (p=0.0597, two-side t student test). The codes of the clones correspond to those indicated in Fig. 1.

We first turned our attention to the T3SS because its inactivation was required to activate symbiosis in the evolution experiment, presumably because some T3SS effectors block nodulation and early infection ^13^. In contrast, the emergence of legume symbiosis *in natura* seems to be associated with the acquisition of T3SS since all rhizobial *Cupriavidus* strains of our sample encode a (chromosomal) T3SS, while most of the other *Cupriavidus* strains do not (Fig. 2). This apparent contradiction is solved by the fact that we could not find a single ortholog of the 77 T3SS effectors of *R. solanacearum* GMI1000 in *C. taiwanensis* LMG19424. Actually, it has been shown that a functional T3SS is not required for mutualistic symbiosis of the latter with *M. pudica* ^24^, the only plant species used in the evolution experiment.

We then focused on PhcA-associated genes since they accumulated an excess of mutations in the experiment (Table S13). The *phc* system, which was only found intact in *Cupriavidus* and *Ralstonia* (Table S14), regulates a reversible switch between two different physiological states via the repression of the central regulator PhcA in *Ralstonia* ^23^ and *Cupriavidus* ^25^. Interestingly, PhcA-associated genes were also enriched in substitutions *in natura*. Indeed, the *phcBQRS* genes of the cell density-sensing system were among the 67 genes that exhibited an excess of nucleotide diversity in the branch bLCA^Ct^ relative to *C. taiwanensis* (“phcA-linked” in Table S4). Strikingly, only seven genes showing an excess of diversity at bLCA^Ct^ had orthologs with mutations in the evolution experiment. Among these seven, only two also showed signature of positive selection in *C. taiwanensis*: *phcB* and *phcS* (ongoing events, Table S3).

Given the parallels between experimental and natural evolution regarding an over-representation of changes in PhcA-associated genes, we enquired on the possibility that mutations in the *phcB*, *phcQ* and *phcS* genes, detected in the evolved E16, K16 and M16 clones capable of nodule cell infection were adaptive for symbiosis with *M. pudica*. For this, we introduced the mutated alleles of these genes in their respective nodulating ancestors, CBM212, CBM349 and CBM356, and the wild-type allele in the evolved clones E16, K16 and M5 (M5 was used instead of M16, since genetic transformation failed in the latter clone in spite of many trials). Competition experiments between the pairs of clones harboring the wild type or the mutant alleles confirmed that these mutations were adaptive (Fig. 4B). The evolved clones also showed better infectivity, since they contained more bacteria per nodule (Fig. S5). On the other hand, we found that the Phc system plays a role in the natural *C. taiwanensis-M. pudica* symbiosis: a *phcA* deletion mutant had lower nodulation competitiveness than the wild-type *C. taiwanensis* (Fig. 4B), and lower infectiveness (Fig. S6), when both strains were co-inoculated to *M. pudica*. Hence, the re-wiring of the *phc* virulence regulatory pathway of *R. solanacearum* was involved in the evolution of symbiosis in several lineages of the experimental evolution. In parallel, high genetic diversification accompanied by positive selection of the homologous pathway was associated with the transition to symbiosis in the natural evolution of *C. taiwanensis*.

## Discussion

Years of comparative genomics and loss of function approaches led to propose that most legume symbionts evolved in two-steps ^8^, *i. e.* acquisition of a set of essential symbiotic genes followed by subsequent adaptation of the resulting genome under plant selection pressure. Although, this evolutionary scenario has recently been validated in the laboratory ^13,17^, to which extent experimental evolution of symbionts parallels natural symbiogenesis was still unknown. Here, we highlighted several parallels between the experimental and *in natura* transitions towards legume symbiosis (Fig. 5). Such parallels were not necessarily expected, because the two processes differed in a number of fundamental points. The two species are from different genera and had different original lifestyles, saprophytic for *C. taiwanensis* and pathogenic for *R. solanacearum*. The conditions of the experimental evolution were extremely simplified and controlled, whereas natural environmental conditions were certainly very complex and changing. The time span of both processes was radically different, 12-16 MYA in nature, and ca. 400 bacterial generations per lineage in the experiment, providing very different magnitudes of genetic diversity. This precluded the identification of parallelisms at the scale of nucleotide positions due to excessive diversity in natural populations. Lastly, *C. taiwanensis* are well-adapted mutualistic symbionts of *Mimosa* spp., whereas the lab-evolution of *Ralstonia* is not yet achieved, none of the evolved clones being able to persist within nodule cells and fix nitrogen to the benefit of the plant.

**Figure 5.**
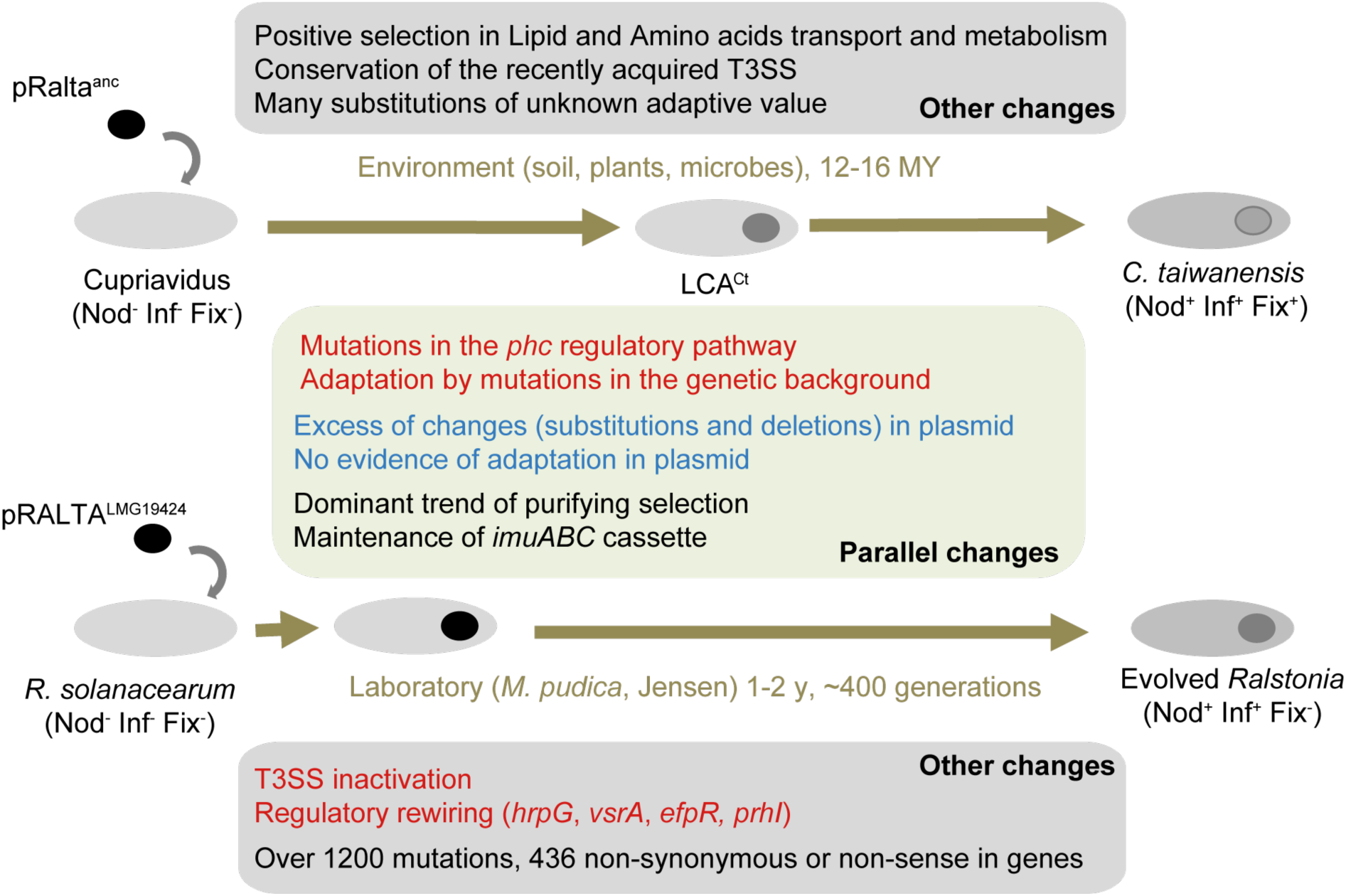
Overall similarities and differences between the experimental and natural evolutionary processes described in this study. Adaptive and non-adaptive changes are in orange and blue, respectively.

The plasmid carrying the essential *nod* and *nif* genes drove the transition towards symbiosis in both processes. We expected that plasmid genes would show evidence of adaptation, either at the level of gene expression regulation or biochemical fine-tuning, to the novel genetic background and environmental conditions. Instead, the abundant substitutions observed in the plasmid seem to have a negligible role in the experiment and lack evidence of positive selection in nature. The cost of the plasmid has also not changed during the experiment. This suggests that the symbiotic genes acquired by *C. taiwanensis* in nature were already - like in the experiment - pre-adapted to establish a symbiotic association with *Mimosa* species. It is in agreement with proposals that pRalta was acquired from *Burkholderia* ^26^, which are ancient symbionts of *Mimosa* spp. ^27^. This also suggests that adaptation following the acquisition of a large plasmid encoding traits driving ecological shifts does not require plasmid evolution. The fact that genetic adaptation to this novel complex trait only occurred in the background is a testimony of the ability of mobile genetic elements to seamlessly plug novel functions in their hosts.

Instead of affecting directly the novel genetic information, adaptive mutations seem to have centered on the rewiring of regulatory modules to inactivate or co-opt native functions for the novel trait. We previously showed that loss of the ability to express the T3SS was strictly necessary for the early transition towards symbiosis in the experiment ^13^, and that subsequent adaptation favored the re-use of regulatory modules leading to massive metabolic and transcriptomic changes ^17^. These phenotypic shifts occurred via mutations targeting regulatory genes specific to *Ralstonia* (*e.g.*, *hrpG, prhI, efpR*, Rsc0965), which finely control the expression of many virulence determinants ^23,28,29^. Here, from the analysis of orthologs between *R. solanacearum* and *C. taiwanensis,* we showed that several genes in the *phcBQRS* operon both exhibited significant positive selection in *C. taiwanensis* populations and accumulated adaptive mutations in the evolution experiment. In *R. solanacearum*, these genes control the activity of the global virulence regulator PhcA via a cell density-dependent mechanism ^30^. Mutations in these genetic regulators are unlikely to cause adaptation by attenuating the virulence of *Ralstonia*, since the chimeric ancestor is not pathogenic on *M. pudica* (and these mutations induced the loss of pathogenicity on *Arabidopsis thaliana* ^18^). Since PhcA also plays a role in the natural *C. taiwanensis-M. pudica* symbiosis, we speculate that adaptive mutations in the experiment and high diversification in nature on *phc* genes after the acquisition of pRalta may reflect the re-wiring of a quorum-sensing system to sense the environment for cues of when to express the novel mutualistic dialogue with eukaryotes. Further work should determine if some of these mutations resulted in the integration of the gene expression network of the plasmid in the broader network of the cell.

Very controlled experimental evolution studies show few similar parallel mutations between replicates and require higher-order analyses at the level of genes, operons or pathways to identify commonalities ^31^. Here, the comparison of the natural evolution of *Mimosa* symbionts in the *Cupriavidus* genus and the experimental symbiotic evolution of *Ralstonia* under *M. pudica* selection pressure could not reveal parallel changes at the nucleotide level because of the high diversity of natural populations. Yet, it showed that symbiotic adaptation occurred in the recipient genome, with similar population genetic patterns, and involved changes in an homologous central regulatory pathway in both processes. These parallels highlight the potential of research projects integrating population genomics, molecular genetics, and evolution experiment to provide insights on adaptation in nature and in the laboratory. Therefore, experimental evolution appears not only useful to demonstrate the biological plausibility of theoretical models in evolutionary biology, but also to enlighten the natural history of complex adaptation processes.

## Methods

### Dataset for the experimental evolution

We used previously published data on the genomic changes observed in the experimental evolution of the chimera, including 21 bacterial clones (three ancestors and 18 evolved clones) ^15^. We analyzed all the synonymous and non-synonymous mutations of each clone from these datasets (Table S12). Large deletions above 1 kb were first listed based on the absence of Illumina reads in these regions, and were then validated by PCR amplification using specific primers listed in Table S15. Primers were designed to amplify either one or several small fragments of the putative deleted regions or the junction of these deletions. All primer pairs were tested on all ancestors and final clones (Table S1).

### Mutant construction

The pRalta in evolved *Ralstonia* clones B16, G16 and I16 or their derivatives, was replaced by the wild-type pRalta of *C. taiwanensis* LMG19424 strain as previously described ^15^, generating B16-op, G16-op and I16-op. Wild-type alleles of the *phcB*, *phcQ* and *phcS* genes and constitutively expressed reporter genes (GFP, mCherry) were introduced into *Ralstonia* evolved clones using the MuGent technique ^32^. Briefly, this technique consisted in the co-transformation of two DNA fragments, one fragment carrying a kanamycin resistance cassette together with a gene coding a fluorophore and one unlabelled PCR fragment of *ca.* 6 kb carrying the point mutation to introduce, as previously described ^17^. Co-transformants were first selected on kanamycin, then screened by PCR for the presence of the point mutation. M5, which possesses the *phcS* mutation, was used instead of M16 since M16 is no more transformable.

To construct the *phcA* deletion mutant of LMG19424, we used the pGPI-SceI/pDAI-SceI technique previously described ^33^. Briefly the regions upstream and downstream *phcA* were amplified with the oCBM3413-3414 and oCBM3415-3416 primer pairs and the Phusion DNA polymerase (Thermo Fisher scientific). The two PCR products were digested with *Xba*I*Bam*HI and *Bam*HI-*Eco*RI respectively and cloned into the pGPI-SceI plasmid digested by *Xba*I and *Eco*RI. The resulting plasmid was introduced into LMG19424 by triparental mating using the pRK2013 as helper plasmid. Deletion mutant were obtained after introduction of the pDAI-SceI plasmid encoding the I-SceI nuclease. LMG19424 *phcA* deletion mutants were verified by PCR using the oCBM3417-3418 and oCBM3419-3420 primer pairs corresponding to external and internal regions of *phcA*, respectively. Oligonucleotides used in these constructions are listed in Table S16.

### Relative *in planta* fitness

*Mimosa pudica* seeds from Australia origin (B &T World Seed, Paguignan, France) were cultivated as previously described ^15^. To measure the *in planta* relative fitness, a mix of two strains bearing different antibiotic resistance genes or fluorophores (5.10^5^ bacteria of each strain per plant) were inoculated to 20 plants. Nodules were harvested 21 days after inoculation, pooled, surface sterilized and crushed. Dilutions of nodule crushes were spread on selective plates, incubated two days at 28°C, then colonies were counted using a fluorescent stereo zoom microscope V16 (Zeiss) when needed. Three independent experiments were performed for each competition.

### Public genome dataset

We collected 13 genomes of *Cupriavidus* spp. (including three rhizobia) and 31 of *Ralstonia* from GenBank RefSeq and the MicroScope platform (http://www.genoscope.cns.fr/agc/microscope/home) as available in September 2015. We removed the genomes that seemed incomplete or of poor quality, notably those smaller than 5 Mb and with L90>150 (defined as the smallest number of contigs whose cumulated length accounts for 90 % of the genome). All accession numbers are given in Table S17. Genomes of α- and β-Proteobacteria larger than 1 Mb and genomes of phages were downloaded from GenBank RefSeq as available in February 2013.

### Sequencing, assembly, and annotation of Illumina data

The genomes of 43 *Mimosa* spp. isolates, a non rhizobial strain of *Cupriavidus* (strain LMG19464) as well as a *C. oxalaticus* strain (LMG2235) (Table S17), were sequenced at the GeT-PlaGe core facility, INRA Toulouse (get.genotoul.fr). DNA-seq libraries were prepared according to Biooscientific’s protocol using the Biooscientific PCR free Library Prep Kit. Briefly, DNA was fragmented by sonication, size selection was performed using CLEANNA CleanPCR beads and adaptators were ligated to be sequenced. Library quality was assessed using an Advanced Analytical Fragment Analyser and libraries were quantified by qPCR using the Kapa Library Quantification Kit. DNA-seq experiments were performed on an Illumina HiSeq2000 sequencer using a paired-end read length of 2 x 100 bp with the HiSeq v3 reagent kit (LMG2235 and LMG19431) or on an Illumina MiSeq sequencer using a paired-end read length of 2 x 300 pb with the Illumina MiSeq v3 reagent kit (other strains). On average, genomes contained 99 contigs and an L90 of 29.

Genome assemblies were performed with the AMALGAM assembly pipeline (Automated MicrobiAL Genome AsseMbler; Cruveiller S. and Séjourné M., unpublished). The pipeline is a python script (v2.7.x and onward) that launches the various parts of the analysis and checks that all tasks are completed without error. To date AMALGAM embeds SPAdes, ABySS ^34^, IDBA-UD ^35^, Canu ^36^, and Newbler ^37^. After the assembly step, an attempt to fill scaffolds/contigs gaps is performed using the gapcloser software from the SOAPdenovo2 package ^38^. Only one gap filling round was performed since launching gapcloser iteratively may lead to an over-correction of the final assembly. AMALGAM ends with the generation of a scaffolds/contigs file (fasta format) and a file describing the assembly in agp format (v2.0).

The genomes were subsequently processed by the MicroScope pipeline for complete structural and functional annotation ^39^. Gene prediction was performed using the AMIGene software ^40^ and the microbial gene finding program Prodigal ^41^ known for its capability to locate the translation initiation site with great accuracy. The RNAmmer ^42^ and tRNAscan-SE ^43^ programs were used to predict rRNA and tRNA-encoding genes, respectively. Genome sequence and annotation was made publicly available (see accession numbers in Table S17).

### PacBio sequencing

Library preparation and sequencing were performed according to the manufacturer’s instructions “Shared protocol-20kb Template Preparation Using BluePippin Size Selection system (15kb-size cutoff)”. At each step DNA was quantified using the Qubit dsDNA HS Assay Kit (Life Technologies). DNA purity was tested using the nanodrop (Thermofisher) and size distribution and degradation assessed using the Fragment analyzer (AATI) High Sensitivity DNA Fragment Analysis Kit. Purification steps were performed using 0.45X AMPure PB beads (Pacbio). 10μg of DNA was purified then sheared at 40kb using the meraruptor system (diagenode). A DNA and END damage repair step was performed on 5μg of sample. Then blunt hairpin adapters were ligated to the library. The library was treated with an exonuclease cocktail to digest unligated DNA fragments. A size selection step using a 13-15kb cutoff was performed on the BluePippin Size Selection system (Sage Science) with the 0.75% agarose cassettes, Marker S1 high Pass 15-20kb.

Conditioned Sequencing Primer V2 was annealed to the size-selected SMRTbell. The annealed library was then bound to the P6-C4 polymerase using a ratio of polymerase to SMRTbell at 10:1. Then after a magnetic bead-loading step (OCPW), SMRTbell libraries were sequenced on RSII instrument at 0.2nM with a 360 min movie. One SMRTcell was used for sequencing each library. Sequencing results were validated and provided by the Integrated next generation sequencing storage and processing environment NG6 accessible in the genomic core facility website ^44^.

### Core genomes

Core genomes were computed using reciprocal best hits (hereafter named RBH), using end-gap free Needleman-Wunsch global alignment, between the proteome of *C. taiwanensis* LMG19424 or *R. solanacearum* GMI1000 (when the previous was not in the subclade) as a pivot (indicated by ** on Fig. S1A) and each of the other 88 proteomes ^45^. Hits with less than 40 % similarity in amino acid sequence or more than a third of difference in protein length were discarded. The lists of orthologs were filtered using positional information. Positional orthologs were defined as RBH adjacent to at least two other pairs of RBH within a neighbourhood of ten genes (five up- and five down-stream). We made several sets of core genomes (see Fig. S1A): all the 89 strains (A1), 44 *C. taiwanensis* (Ct), Ct with the closest outgroup (C2), Ct with the five closest outgroups (C3), the whole 60 genomes of the genus *Cupriavidus* (Cg), and the 14 genomes of *R. solanacearum* (Rs). They were defined as the intersection of the lists of positional orthologs between the relevant pairs of genomes and the pivot (Table S18).

### Pan genomes

Pan genomes describe the full complement of genes in a clade and were computed by clustering homologous proteins in gene families. Putative homologs between pairs of genomes were determined with blastp v2.2.18 (80 % coverage), and evalues (if smaller than 10^-4^) were used to infer protein families using SiLiX (v1.2.8, http://lbbe.univlyon1.fr/SiLiX) ^46^. To decrease the number of paralogs in pan genomes, we defined a minimal identity threshold between homologs for each set. For this, we built the distribution of identities for the positional orthologs of core genomes between the pivot and the most distant genome in the set (Fig. S7), and defined an appropriate threshold in order to include nearly all core genes but few paralogs (Table S19).

### Alignment and phylogenetic analyses

Multiple alignments were performed on protein sequences using Muscle v3.8.31 ^47^, and back-translated to DNA. We analyzed how the concatenated alignment of core genes fitted different models of protein or DNA evolution using IQ-TREE v1.3.8 ^48^. The best model was determined using the Bayesian information criterion (BIC). Maximum likelihood trees were then computed with IQ-TREE v1.3.8 using the appropriate model, and validated via a ultrafast bootstrap procedure with 1000 replicates ^49^ (Table S18). The maximum likelihood trees of each set of core genes were computed with IQ-TREE v1.3.8 using the best model obtained for the concatenated multiple alignment.

In order to root the phylogeny based on core genes, we first built a tree using 16S rRNA sequences of the genomes of *Ralstonia* and *Cupriavidus* genera analysed in this study and of ten outgroup genomes of β-Proteobacteria. For this, we made a multiple alignment of the 16S sequences with INFERNAL v.1.1 (with default parameter) ^50^ using RF00177 Rfam model (v.12.1) ^51^, followed by manual correction with SEAVIEW to removed poorly aligned regions. The tree was computed by maximum likelihood with IQ-TREE using the best model (GTR+I+G4), and validated via an ultrafast bootstrap procedure with 1000 replicates.

To date the acquisition of the symbiotic plasmid in the branch bLCA^Ct^, we computed the distances in the 16S rDNA tree between each strain and each of the nodes delimitating the branch bLCA^Ct^ (respectively LCA^Ct^ and C2 in Fig. 2). The substitution rate of 16S in enterobacteria was estimated at ∼1% per 50 MY of divergence ^52^, and we used this value as a reference.

### Orthologs and pseudogenes of symbiotic genes, the mutagenic cassette, T3SS and PhcABQRS

We identified the positional orthologs of Cg for symbiotic genes, the mutagenic cassette, T3SS, and PhcABQRS using RBH and *C. taiwanensis* LMG19424 as a pivot (such as defined above). These analyses identify *bona fide* orthologs in most cases (especially within species), and provide a solid basis for phylogenetic analyses. However, they may miss genes that evolve fast, change location following genome rearrangements, or that are affected by sequence assembling (incomplete genes, small contigs without gene context, etc.). They also miss pseudogenes. Hence, we used a complementary approach to analyze in detail the genes of the symbiotic island in the plasmid, the mutagenic cassette, T3SS and PhcABQRS. Indeed, we searched for homologs of each gene in the reference genome in the other genomes using LAST v744 ^53^ and a score penalty of 15 for frameshifts. We discarded hits with evalues below 10^-5^, with less than 40 % similarity in sequence, or aligning less than 50 % of the query. In order to remove most paralogs, we plotted values of similarity and patristic distances between the 59 *Cupriavidus* and the reference strain *C. taiwanensis* LMG19424 for each gene. We then manually refined the annotation using this analysis.

### Evolution of gene families

We used Count (version downloaded in December 2015) ^54^ to study the past history of transfer, loss and duplication of the protein families of the pan genomes. The analysis was done using the core genomes reference phylogenies. We tested different models of gene content evolution using the tree of Cg (Table S18), and selected the best model using the Akaike information criterion (AIC) (Table S19). We computed the posterior probabilities for the state of the gene family repertoire at inner nodes with maximum likelihood and used a probability cutoff of 0.5 to infer the dynamics of gene families, notably presence, gain, loss, reduction, and expansion for the branch leading to the last common ancestor (LCA) of *C. taiwanensis* (LCA^Ct^).

### Measures of similarity between genomes

For each pair of genomes, we computed two measures of similarity, one based on gene repertoires and another based on the sequence similarity between two genomes. The gene repertoire relatedness (GRR) was computed as the number of positional orthologs shared by two genomes divided by the number of genes in the smallest one ^55^. Pairwise average nucleotide identities (ANIb) were calculated using the pyani Python3 module (https://github.com/widdowquinn/pyani), with default parameters ^56^. We used single-linkage clustering to group strains likely to belong to the same species. This was done constructing a transitive closure of sequences with an ANIb higher than a particular threshold (*i.e.*, >94%, 95% or 96%). We used BioLayout Express^3D^ to visualize the graphs representing the ANIb relationships and the resulting groups for each threshold (Fig. S8).

### Inference of recombination

We identified recombination events using three different approaches. We used the pairwise homoplasy index (PHI) test to look for incongruence within each core gene multiple alignment (Ct and C3 datasets). We made 10,000 permutations to assess the statistical significance of the results ^57^. We used the SH-test, as implemented in IQ-TREE v1.3.8 ^48^ (GTR+I+G4 model, 1000 RELL replicates), to identify incongruence between the trees of each core gene and the concatenated multiple alignment of all core genes. We used ClonalFrameML v10.7.5 ^58^ to infer recombination and mutational events in the branch leading to the LCA^Ct^ using the phylogenetic tree of C3 (Table S18). The transition/transversion ratios given as a parameter to ClonalFrameML were estimated with the R package PopGenome v2.1.6 ^59^. Lastly, ClonalFrameML was also used to compare the relative frequency of recombination and mutation on the whole concatenated alignments of Ct and Rs.

### Molecular diversity and adaptation

Positive selection was identified using likelihood ratio tests by comparing the M7 (beta) - M8 (beta &ω) models of codeml using PAML v4.8 ^60^. We used the independent phylogenetic tree of each gene family to avoid problems associated with horizontal transfer (since many genes failed the SH-test for congruence with the core genome phylogenetic tree). We removed from the analysis gene families that had incongruent phylogenetic signals within the multiple alignment ^61^. These correspond to the families for which PHI identified evidence of recombination (p < 0.05).

We inferred the mutations arising in the branch leading to LCA^Ct^ using the phylogenetic tree build with the core genome of C3 (Ct and the five closest outgroups). First, we used ClonalFrameML to reconstruct the ancestral sequences of LCA^Ct^ and LCA^C2^ (accounting for recombination). Then, we estimated nucleotide diversity of each core gene for Ct, and between LCA^Ct^ and LCA^C2^ using the R package pegas. Finally, we used the branch-site model of codeml to identify positive selection on this branch for the core genes of C3 that lacked evidence of intragenic recombination (detected using PHI).

To infer the extent of purifying selection for Ct, we computed dN/dS values for each core genes between *C. taiwanensis* LMG19424 and the others strains of Ct using the yn00 model of PAML v4.8. We then plotted the average dN/dS of each strains with the patristic distances obtained from the tree of the concatenated multiple alignment of all core genes.

### Functional annotations

We searched for the functions over-represented relative to a number of characteristics (recombination, nucleotide diversity, etc.). We analyzed COG categories, protein localizations, transporters, regulatory proteins and several pre-defined lists of genes of interest in relation to rhizobial symbiosis and to virulence.

We used COGnitor ^62^ as available on the MicroScope Platform (https://www.genoscope.cns.fr/agc/microscope/home) to class genes according to the COG categories (Tables S3 and S8). Protein subcellular localizations were predicted using PSORTb v3.0.2 (http://www.psort.org/psortb) ^63^. Transporters and regulatory proteins were inferred using TransportDB (http://www.membranetransport.org) ^64^ and P2RP (http://www.p2rp.org) ^65^, respectively. Protein secretion systems were identified using TXSScan (http://mobyle.pasteur.fr/cgi-bin/portal.py#forms::txsscan) ^66^. We manually checked and corrected the lists. Specific annotations were also defined for (i) *R. solanacearum* GMI1000: Type III effectors ^67^, PhcA-associated genes (*i.e.*, genes involved in the upstream regulatory cascade controlling the expression of phcA, and genes directly controlled by PhcA) ^23,68,69^, virulence ^70^, extracellular polysaccharides (EPS) ^69,71^, chemotaxis ^72^, twin-arginine translocation pathway (Tat) ^73^, Tat-secreted protein ^74^, and (ii) the pRalta of *C. taiwanensis* LMG19424: symbiotic genes ^14^, genes pertaining to plasmid biology (conjugation, replication, partition, based on the annotations ^75^), and operons using ProOpDB (http://operons.ibt.unam.mx/OperonPredictor) ^76^. Lastly, we also annotated positional orthologs between *R. solanacearum* GMI1000 and *C. taiwanensis* LMG19424 according to specific annotations used for both strains (Tables S3 and S8).

### Analysis of the mutations observed in the experimental evolution

To estimate differences between mutation rates on the three replicons of the chimera, we compared the observed number of synonymous mutations in each replicon to those obtained from simulations of genome evolution. First, we analyzed the distribution of synonymous mutations of the 18 final evolved clones in regions of the genome that were covered by sequencing data (some regions with repeats cannot be analyzed without ambiguity in the assignment of mutations). We built the mutation spectrum of the genome using these synonymous mutations, since they are expected to be the least affected by selection. Second, we performed 999 random experiments of genome evolution using the mutation spectrum and the total number of synonymous mutations obtained for the 18 final clones. With the results, we draw the distributions of the expected number of synonymous mutations in each replicon (under the null hypothesis that they occurred randomly). This data was then used to define intervals of confidence around the average values observed in the simulations.

### Statistical analyses

In order to identify genes that evolved faster in the branch leading to LCA^Ct^, we compared the nucleotide diversity of sequences for LCA^Ct^ and LCA^C2^ with those of the extant 44 *C. taiwanensis* using a regression analysis. Outliers above the regression line were identified using a one-sided prediction interval (p < 0.001) as implemented in JMP (JMP^®^, Version 10. SAS Institute Inc., Cary, NC, 1989-2007).

We computed functional enrichment analyses to identify categories over-represented in a focal set relative to a reference dataset. The categories that were used are listed above in the section *Functional annotations*. To account for the association of certain genes to multiple functional categories, enrichments were assessed by resampling without replacement the appropriated reference dataset (see Table S20) to draw out the expected null distribution for each category. More precisely, we made 999 random samples of the number of genes obtained for each analysis (positive selection, recombination, etc.) in the reference dataset. For each category, we then compared the observed value (in the focal set) to the expected distribution (in the reference dataset) to compute a p-value based on the number of random samples of the reference dataset that showed higher number of genes from the category.

We also compared the nucleotide diversity between sets of genes using the nonparametric Wilcoxon rank sum test ({stats}, wilcox.test).

Finally, we computed Fisher’s exact tests (R package {stats}, fisher.test) to estimate the association between results of the natural and the experimental evolution, *i.e.*, to test whether mutations found in the experimental evolution targeted genes that were found to be significantly more diverse in the natural process.

P-values were corrected for multiple comparisons using Benjamini and Hochberg’s method ^77^ ({stats}, p.adjust).

Statistical analyses with R were done using version 3.1.3 (R: a language and environment for statistical computing, 2008; R Development Core Team, R Foundation for Statistical Computing, Vienna, Austria [http://www.R-project.org]).

## Acknowledgements

This work was supported by funds from the French National research Agency (ANR-12-ADAP-0014-01 and ANR-16-CE20-0011-01) the “Laboratoire d’Excellence (LABEX)” TULIP (ANR-10-LABX-41) and France Génomique National infrastructure, funded as part of “Investissement d’avenir” program managed by Agence Nationale pour la Recherche (contrat ANR-10-INBS-09).

We thank Olaya Rendueles, Rémi Peyraud, Ludovic Cottret, Stéphane Genin, and Pedro Couto Oliveira for helpful comments and suggestions. We thank Eddy Ngonkeu and Moussa Diabate for help with the strain collection.

## Contributions

CC, CM, and ER conceived the project, integrated the analyses, and wrote the draft of the manuscript. CC, MTouchon, and ER made the computational analyses. DC and MTang performed the experiments and analyzed the data. LM and MAP provided strains and data. SC assembled and annotated the genomes. All authors contributed to the final text.

